# Flavor compounds in fermented Chinese alcoholic beverage alter gut microbiota and attenuate ethanol-induced liver damages

**DOI:** 10.1101/437731

**Authors:** Cheng Fang, Hai Du, Zheng Xiaojiao, Aihua Zhao, Wei Jia, Yan Xu

## Abstract

Alcoholic beverages which are consumed widely in most parts of the world have long been identified as a major risk factor for all liver diseases, particularly alcohol-induced liver disease (ALD). Recent compositional analyses suggest that Chinese Baijiu (CB), a clear alcoholic liquid distilled from fermented grains, contains large amounts of small molecule bioactive compounds in addition to a significant amount of ethanol. Here, in an experimental mouse model, we show that CB caused lower degrees of liver injury than pure ethanol by protecting against the decrease of the relative abundance of *Akkermansia* and increased relative abundance of *Prevotella* in the gut thereby preventing the destruction of the intestinal barrier. Furthermore, we demonstrated that ethanol-induced alteration of the gut microbiota profoundly affected the host metabolome. Compared with ethanol feeding, CB feeding resulted in higher concentrations of functional saturated LCFAs and SCFAs. Our results provide supporting evidence that ALD was profoundly influenced by host-gut microbiota metabolic interactions and that small molecule organic compounds in CB could attenuate ALD.

## Introduction

As a public health problem, alcoholic liver disease (ALD) is a major factor in the burden of disease and also one of the leading causes of liver-related deaths worldwide^1,2^ A recent epidemiological study has estimated that approximately 50% of all cirrhosis-associated deaths are related to alcohol^3^. Hence, the health and social issues related to alcohol use and misuse need attention urgently. However, a growing number of researchers have suggested that moderate drinking benefits healthy adults in many ways. Drinking in moderation has been reported to reduce the risk of cardiovascular diseases^4^, diabetes^5^, and may have protective effects against ischemic stroke^6^ and osteoporosis^7^.

Although the underlying mechanisms are incompletely understood, the importance of the intestinal microbiota in the pathogenesis of ALD is increasingly recognized^8, 9, 10^. Alcohol-induced gut microbiota dysbiosis destroys the intestinal barrier function and increases the intestinal permeability, which results in increased translocation of bacterial components, bacteria, and metabolites from the gut to the liver through the portal vein and the systemic circulation. Lipopolysaccharide (LPS), a Gram-negative bacteria cell wall component, engages in a complex signaling cascade through different receptors in the innate immune system. By activating the NF-κB pathway, LPS induces transcription of various pro-inflammatory cytokines and chemokines which are considered important mediators of liver-gut communication^11, 12^.

In addition to inducing dysbiosis, alcohol alters the metabolic composition of the gastrointestinal contents^13^. The intestinal microbiome performs a diverse range of metabolic functions including production of numerous metabolites that serve as the nutritional sources for microbes and important messengers between the microbiota and the host^14^. The alteration of the metabolome caused by dysbiosis can find evidence in many metabolic diseases and therefore, this microbiome-metabolome homeostasis is important for the host health^15, 16^. However, supporting scientific data about the microbiome-metabolome interactions remain sparse in mouse models of ALD.

Distilled liquor is a group of important beverage products both culturally and economically. It is a complex mixture of water, ethanol, and thousands of small molecule chemical compounds^17, 18^. Recent studies have shown that many distilled liquors contain bioactive compounds and have certain antioxidant activity. For example, brandy and whiskey were found to have antioxidant activity and this activity was associated with their total phenolic content^19^. Likewise, Chinese Baijiu (CB), a traditional Chinese fermented beverage, was also found to have many compounds with potential bioactivity^20^. CB is produced from multi-strain and solid-state fermentation techniques that produce large numbers of various fermentation metabolites, such as esters, alcohols, aldoketones, acids and many other compounds that were bioactive including short chain fatty acids, phenols and heterocyclic compounds^17^. Therefore, we hypothesized that, in chronic alcoholism, grain-derived fermentation metabolites in CB have a hepatoprotective effect and CB ingestion may cause lower levels of liver injury compared with the same dosage of ethanol (EtOH).

In this study, to verify our hypothesis, we developed a chronic EtOH gavage mouse model and compared the effects of CB and EtOH intervention on the liver. In addition, to elucidate how CB may affect the gut microbiota and their metabolome, and discriminate such changes from EtOH-induced alterations, we compared the effects of CB and EtOH on the intestinal microbiota and metabolic parameters, including short chain fatty acids (SCFAs). As far as we know, this is the first study that reveals the effect of distilled liquor on intestinal microflora and their metabolism. Our results may improve our understanding of the host-gut microbiota metabolic interactions during the progression of ALD.

## Results

### Chinese Baijiu and its compositions

CB used in this study is a distilled liquor produced at Guizhou, China and was purchased from local market. The chemical compositions of CB were identified using a mass spectrometry-based metabolomics approach and provided in Table S1.

### CB and EtOH intervention lowered the food consumption and body weight and increased the liver weight/body weight (LIV/BW) ratio

To explore the effect of EtOH consumption on phenotype, we performed a 12-week intragastric administration experiment. Figure 1A is a diagrammatic representation of the alcohol dosing experiment. During the feeding trial period, all mice were eating and drinking normally except for one EtOH treated mouse which died at week 11. The food consumption of both CB- and EtOH-fed mice, however, were significantly lower relative to control mice (Fig. 1B) and these differences in food consumption were directly reflected in the body weights of the mice (Fig. 1C). The body weight of experimental mice were significantly lower than control mice except for the 8-week CB-fed mice where no significant change was observed (Fig. 1D) based on the fact that the initial weight of mice in each group had no significant differences (Table S2). As expected, the LIV/BW ratio of experimental mice were higher than the control mice and the LIV/BW ratio of EtOH-fed mice were higher than those treated with CB (Fig. 1D), although this difference was observed to be significant only at week 4. The observed increase in the LIV/BW ratio of ethanol-fed mice (either CB or EtOH) led us to hypothesize that the changes were caused by liver injury and fatty liver. The attenuated effect of CB relative to EtOH on the LIV/BW ratio was supporting evidence for CB being less hepatotoxic.

**Figure 1.**
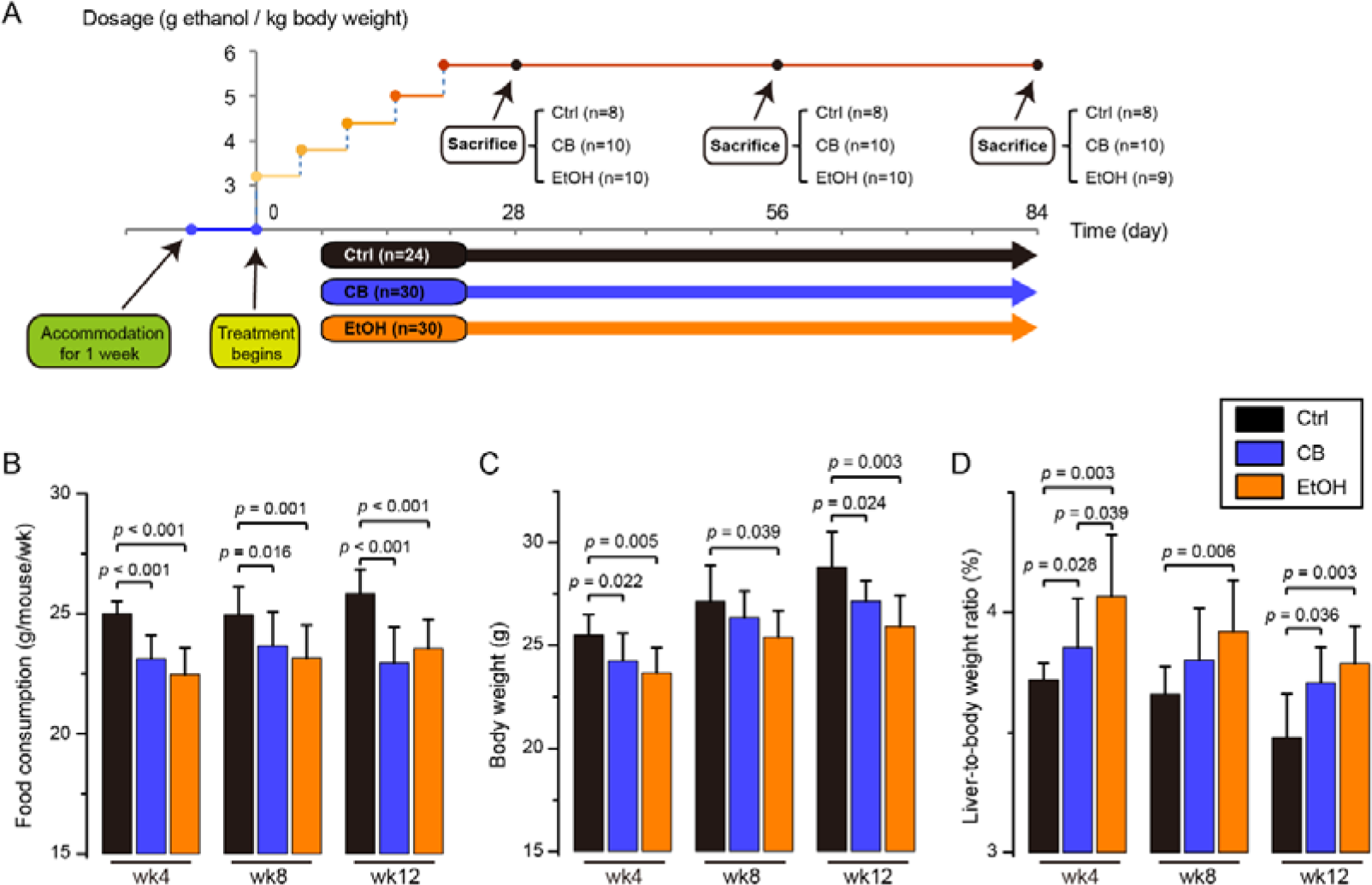
Effect of Chinese Baijiu (CB) and ethanol (EtOH) intervention on mouse phenotype. (A) Animal feeding schedule: mice of intervention groups were given CB (n=30; 10 replicates for each time point) or EtOH (n=29; 9-10 replicates for each time point) by gavage and pair-fed controls (Ctrl, n=24; 8 replicates for each time point) received the vehicle (water). The final dosage (5.7 g ethanol/kg bodyweight) is equivalent to approximately 3 standard human drinks. (B) Food consumption. (C) Body weight and (D) Liver-to-body weight ratio. The Liver-to-body weight ratio (%) was calculated as g/100 g body weight. Bar plot is shown as mean ± S.D. Significance was evaluated using the two-tailed unpaired Student *t*-test.

### CB feeding resulted in a lower level of hepatic injury and steatosis than EtOH feeding

To determine whether different phenotypes caused by CB and EtOH intervention directly translated into differences in ALD, the biochemical and pathological changes induced by CB and EtOH were further characterized. First, steatosis and lipid accumulation in liver tissue were observed by H&E (Figure 2A) and oil red O staining (Fig. 2B). The EtOH-fed mice exhibited both micro- and macrovesicular steatosis at 4 weeks following alcohol administration while it was not observed in CB-fed mice until week 12 (Fig. 2A). Hepatic fat accumulation was also markedly lower in CB-fed mice compared with EtOH-fed mice (Fig. 2B). Biochemical analyses including plasma alanine aminotransferase (ALT) and hepatic triglyceride (TG) levels were measured and although both CB and EtOH feeding lead to significant increases in plasma ALT (Fig. 2C) and accumulation of hepatic TGs (Fig. 2D), CB gavage partially prevented liver injury, as assessed by lower plasma ALT levels, and less TGs relative to EtOH treated mice. In addition, CB-fed mice had significantly lower levels of thiobarbituric acid-reactive substances (TBARS) compared EtOH treated mice which indicated that CB treatment caused lower levels of hepatic oxidative stress (Fig. 2E). These results led us to hypothesize that the bioactive compounds in CB exerted a hepatoprotective effect because even though both CB and EtOH-fed mice were exposed to an equivalent amount of ethanol, there was less EtOH induced toxicity found in the CB group.

**Figure 2.**
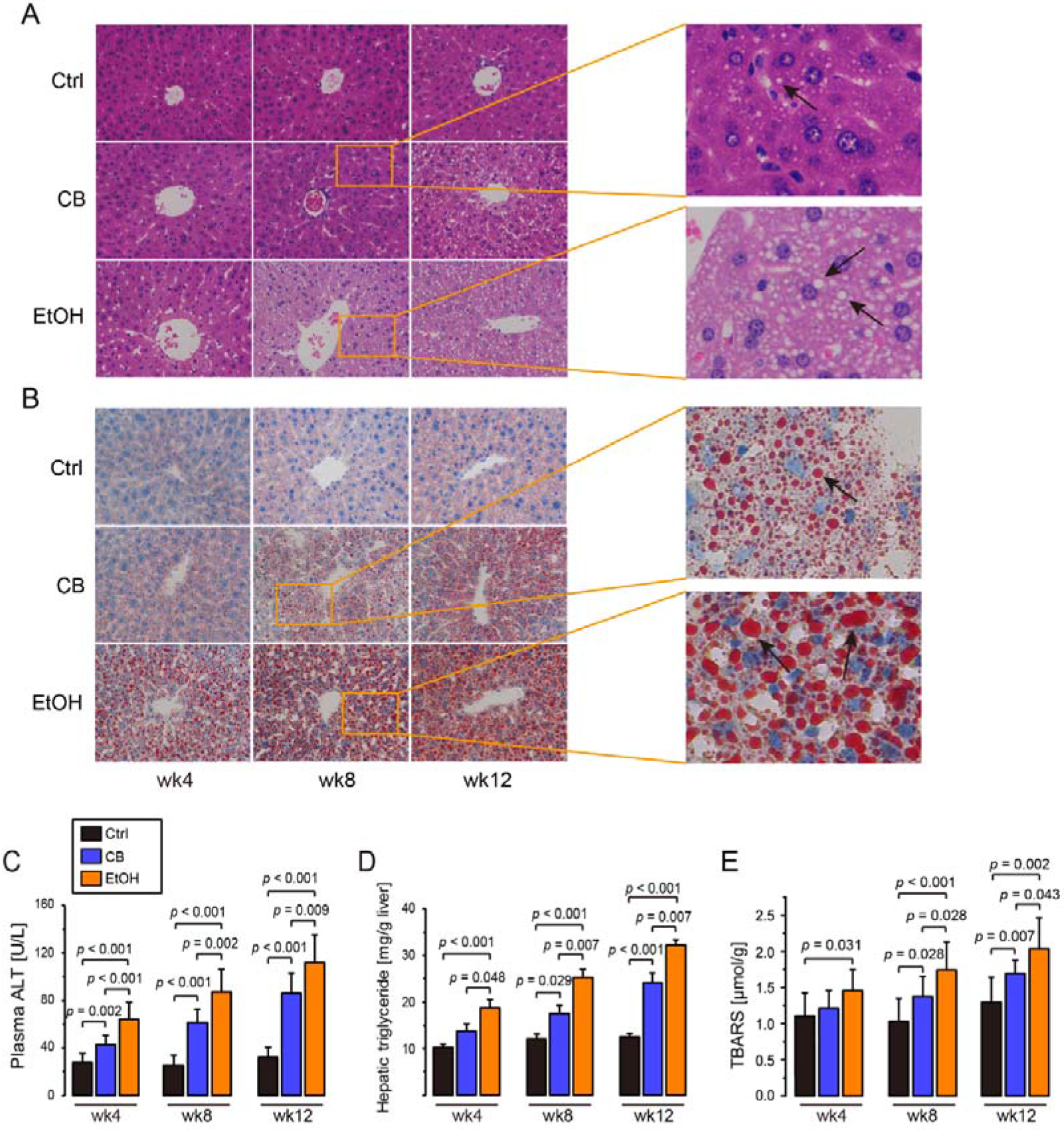
CB caused lower levels of liver injury than EtOH. (A) Representative photomicrographs of H&E liver sections. (B) Representative photomicrographs of Oil Red O-stained liver sections (C) Levels of plasma alanine aminotransferase (ALT) (D) Levels of hepatic triglyceride. (E) Hepatic levels of thiobarbituric acid-reactive substances, Magnification, 400* (n=8 mice/time point for the Ctrl group; n=9-10 mice/time point for the experimental groups). Bar plot is shown as mean ± SEM. Significance was evaluated using the two-tailed unpaired Student *t*-test.

### CB and EtOH feeding resulted in different intestinal community structures

Recent advances in ALD research have elucidated the importance of gut microbiota in the development and progression of ALD^11, 21^. Our findings indicated that CB feeding caused lower levels of liver injury than EtOH feeding. To investigate whether the observed differences in liver injury between mice fed CB and those fed EtOH in our model were associated with the differences in the intestinal microbiota, we analyzed the caecal microbiota by sequencing the V3-V4 amplicons of 16S rRNA genes. We hypothesized that EtOH-fed mice would experience a distinct composition from both control and CB mice.

After trimming, assembly and quality filtering, the resulting sequences were delineated into 657 operational taxonomic units (OTUs) at the similarity cutoff of 97%. Ninety nine percent of OTUs were detected by at least two DNA reads, demonstrating thorough sampling of the gut microbiota. Rarefaction analysis showed that all the caecal microbial diversity in each sample was captured with the current sequencing depth (Fig. S1). OTU-based unweighted unifrac PCoA analysis revealed that the gut microbiota structure of the treated groups showed a time-dependent variation (Fig. 3A) and that the experimental groups significantly differed from the control (Fig. 3B). Taxonomy-based analysis at the phylum level revealed that the phyla Firmicutes, Verrucomicrobia, Proteobacteria and Bacteroidetes dominated the caecal microbial communities (Fig. S2). Experimental mice had notable increases in the phylum Proteobacteria and decreases in phylum Actinobacteria. Although the relative abundances of the phyla Firmicutes and Bacteroidetes showed no significant changes, their ratio, considered to be of significant relevance to the gut microbiota composition^22^, was significantly lower in the treated mice than in the control mice especially for the EtOH-fed mice (Fig. S3). In addition, we also observed that the percentages of Gram-negative (G-) bacteria increased in the experimental groups (Fig. S4). Despite EtOH’s key role in affecting gut flora, clear differences in the bacterial composition between mice fed CB and those fed EtOH were observed and these differences tended to be expanded (Figs. S5, S6). Our results were congruent with multiple lines of evidence that EtOH intake induces gut dysbiosis^23, 24^ and, moreover, CB and EtOH feeding lead to different caecal community structures, thus proving our initial hypothesis that the EtOH-fed mice would have a distinct microbiota composition from that of CB-fed mice.

**Figure 3.**
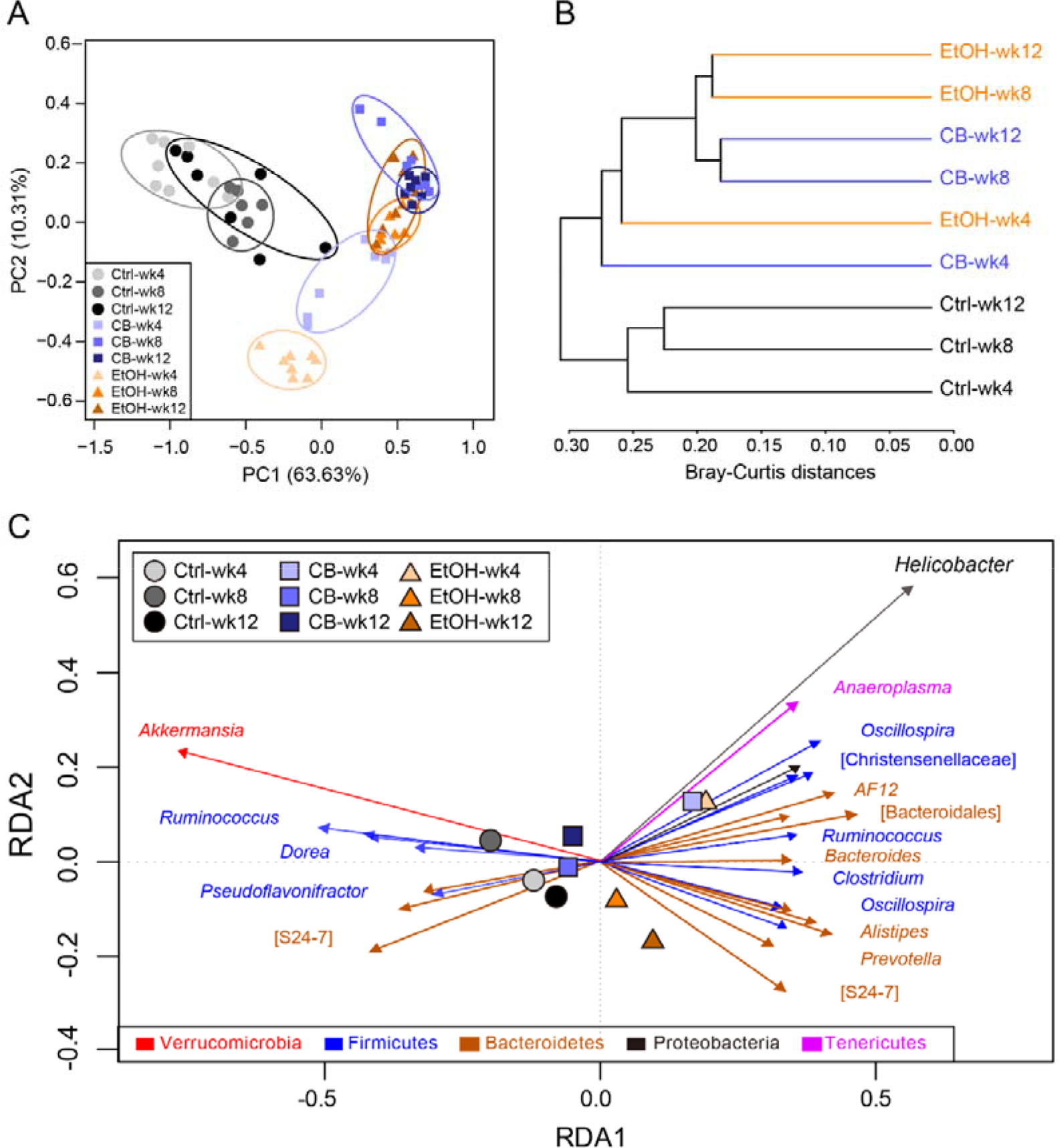
Time-dependent alterations of the gut microbiota and the key OTUs responding to CB or EtOH. (A) Unweighted Unifrac PCoA of gut microbiota based on the OUT data (n=8, 6 and 7 in control group at week 4, 8 and 12, respectively; n=7, 7 and 7 in CB group at week 4, 8 and 12, respectively; n=8, 6 and 7 in EtOH group at week 4, 8 and 12, respectively); (B) Clustering of gut microbiota based on Bray-Curtis distances calculated with multivariate analysis of variance (MANOVA); (C) Biplot of redundancy analysis (RDA) of the caecal microbiota composition responding to CB and EtOH. Different treatments at different times were used as environmental variables. Responding OTUs that explain more than 4% of the sample variability are indicated by arrows. Arrows pointing to the right represented the key OTUs increased by EtOH. Arrows pointing to the left represented the key OTUs decreased by EtOH.

In order to determine the key OTUs responding to the CB and EtOH, redundancy analysis (RDA) was performed using different treatments at different times as environmental variables (Fig. 3C). Similar to the results discussed above, a difference in composition of caecal microbiota between CB- and EtOH-fed mice was observed. Eight- and twelve-week treatments of CB groups are located at the left of the RDA1 together with the control groups, while the 4 week CB treatment group and EtOH groups are located at right of the RDA1. This result further confirmed the differences in the caecal microbial communities between CB and EtOH treatment groups and revealed that the community structure tended to be more similar between CB and control group than between EtOH and control group. In addition, a total of 28 key OTUs were identified, among which 19 OTUs were increased by EtOH. These increased OTUs belong to *Oscillospira* (n=4), *Helicobacter* (n=2), unclassified S24-7 (n=4), and one OTU to each of the following genera: *Ruminococcus, AF12, Alistipes, Clostridium Bacteroides, Prevotella, Anaeroplasma,* unclassified OTUs of the families Bacteroidales and Christensenellaceae. The rest of the 9 OTUs that were decreased by EtOH treatment belonged to the genera *Akkermansia* (n=2), unclassified S24-7 (n=3), *Pseudoflavonifractor* (n=2), *Dorea* (n=1) and *Ruminococcus* (n=1) (Tables S3, S4).

### Intestinal barrier functions of CB-treated mice were more complete relative to EtOH-treated mice

By comparing the key OTUs derived from RDA between CB and EtOH treated groups (Fig. S7), we found that CB-treated mice had higher levels of OTUs affiliated with the genus *Akkermansia* (OTU1) than EtOH-treated mice from week 8 onwards (Fig. 4A). An increasing amount of research has disclosed that the abundance of *Akkermansia* is positively correlated with the integrity of the intestinal barrier^25^. Epithelial tight junctions are critically important to maintain barrier integrity and influence intestinal epithelial leakage^26^. Hence, we hypothesized that a diminished abundance of *Akkermansia* in EtOH-fed mice would result in a decrease in the tight junction protein expression in the ileum tissue of these mice. Although both CB and EtOH treatment reduced the expression of occludin, CB-fed mice had significantly higher levels of occludin than those of EtOH-fed mice (Figs. 4B, C). Claudin-2 forms pores to increase barrier leakiness^27^. Levels of claudin-2 increased in EtOH-fed mice relative to CB-fed mice (Figs. 4D, E). Destruction of intestinal barrier function would lead to leakage of microorganisms and their products and induce endotoxemia. Accordingly, we measured the serum LPS level. As expected, the CB-fed mice had significantly lower levels of serum LPS than EtOH-fed mice (Fig. 4F). These results suggest that CB feeding may attenuate the ethanol-induced decrease of *Akkermansia* populations which, in turn, if at higher abundance, may ameliorate the destruction of intestinal barrier function.

**Figure 4.**
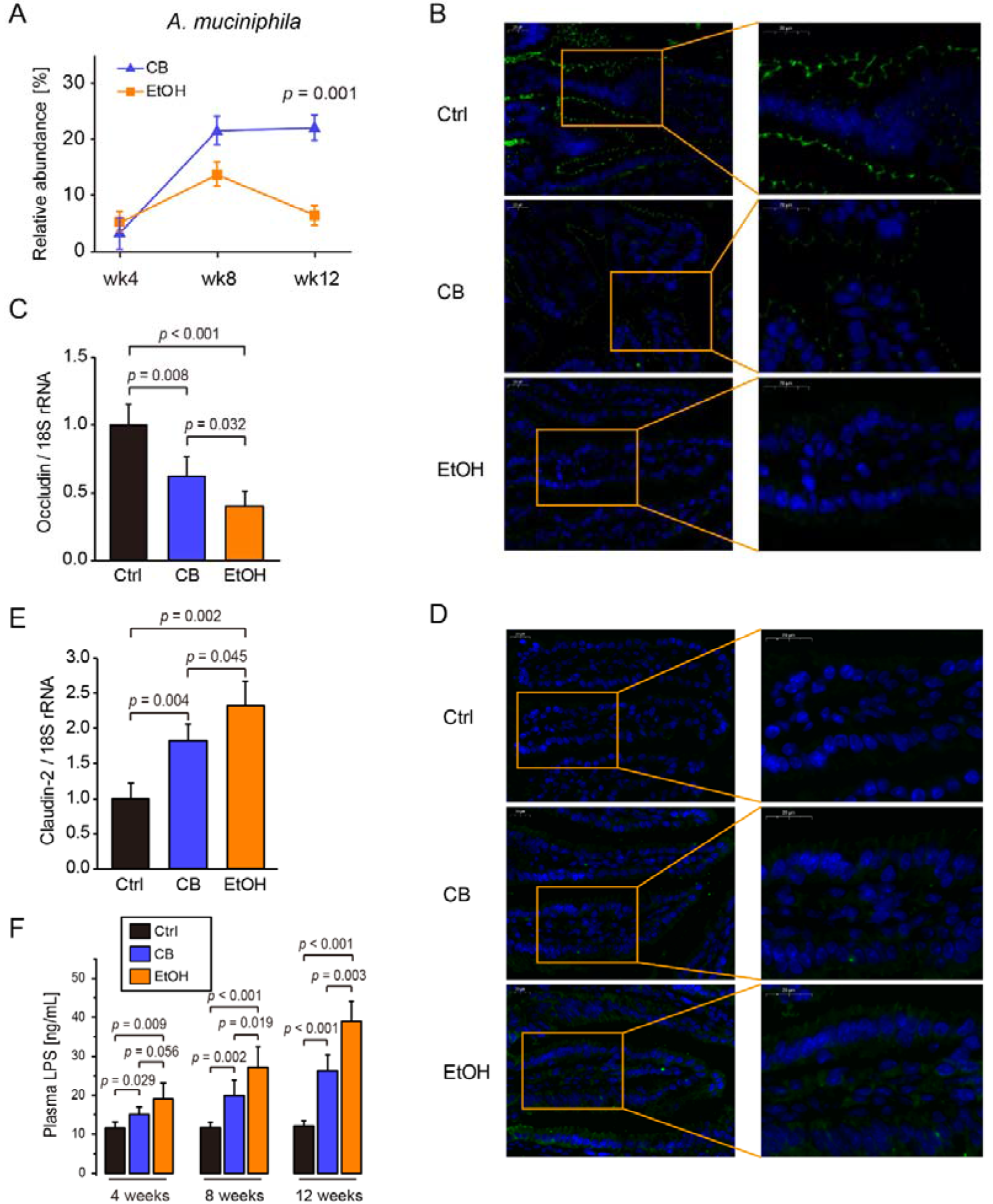
Intestinal barrier functions of CB-fed mice was more complete than EtOH-fed mice (A) Abundance of *A. muciniphila* in cecum responding to CB and EtOH (B) Representative images of immunofluorescence analysis of occludin (green) in distal small intestine of 12-week mice. Nuclei were stained with Hoechst (blue), 400x magnification (C) Quantification of occludin protein normalized to 18S rRNA. (D) Representative images of immunofluorescence analysis of claudin-2 (green) in distal small intestine of 12-week mice. Nuclei were stained with Hoechst (blue), 400* magnification. (E) Quantification of claudin-2 protein normalized to 18S rRNA. (F) LPS level in plasma. Bar plot is shown as mean ± SEM. Significance was evaluated using the two-tailed unpaired Student *t*-test.

### CB and EtOH treatment resulted in two distinct caecal metabolomes

Although the associations between gut microbiota and pathogenesis of ALD has been well established^11, 28^, the effects of the altered gut microbiota on their metabolic profiles are poorly understood. Therefore, in addition to understanding how EtOH and CB altered gut community structure, we investigated how gut community structure impacts function. The metabolic profiles of caecal contents were therefore analyzed using untargeted metabolomics.

The time-dependent metabolic “footprints” of the caecal metabolome showed that control groups were clearly separated from the experimental groups using first principal component (PC1), while the second principal component (PC2) further differentiated the EtOH and CB groups revealing that the distance between them tended to expand in a time-dependent manner (Figs. 5A, B). These results suggested that the altered gut microbiota caused functional changes. To further characterize the metabolic perturbation in response to CB and EtOH feeding, metabolites that were altered were subjected to a two-tailed unpaired *t*-test and statistical significance was decided on the basis of a *p* value of less than 0.05 and a VIP value more than 1. A total of 44 differential metabolites were identified in this way. Among these metabolites, 15 were related to amino acid metabolism, 14 were lipid-related metabolites that were primarily saturated long-chain fatty acid (LCFAs). In addition, there were 8 carbohydrate-related metabolites, 2 that were relevant to gut microbial metabolism, 1 for bile acid metabolism and 4 metabolites were categorized as miscellaneous (Fig. S8). The alteration of metabolites with various functions and our previous data regarding changes in gut microbiota composition suggest that these important impacts were caused by administration of EtOH. Of these differential metabolites, we observed that CB-fed mice had lower concentrations of carbohydrates than that of the EtOH-fed group. Because gut bacterial species across many taxa share the genes for fermenting carbohydrates into short-chain fatty acids (SCFAs)^29^, we hypothesized that CB-fed mice had a higher content of SCFAs in their gut lumen relative to EtOH-fed mice. Hence, we quantified SCFA concentrations in the mouse colonic contents (Fig. 5C). Although both CB and EtOH feeding led to increased acetate and decreased butyrate levels^13^, consistent with our hypothesis, CB-fed mice had higher colonic concentrations of most SCFAs (ie. acetic acid, butyric acid, isobutyric acid and valeric acid) than was found for EtOH-fed mice, especially for acetic and butyric acids (Fig. 5D, E, S9). Similar to the findings for the gut microbiome, these results indicated that although alcohol feeding notably altered the gut metabolome, clear differences in the gut metabolome were observed between mice fed CB and mice fed EtOH.

**Figure 5.**
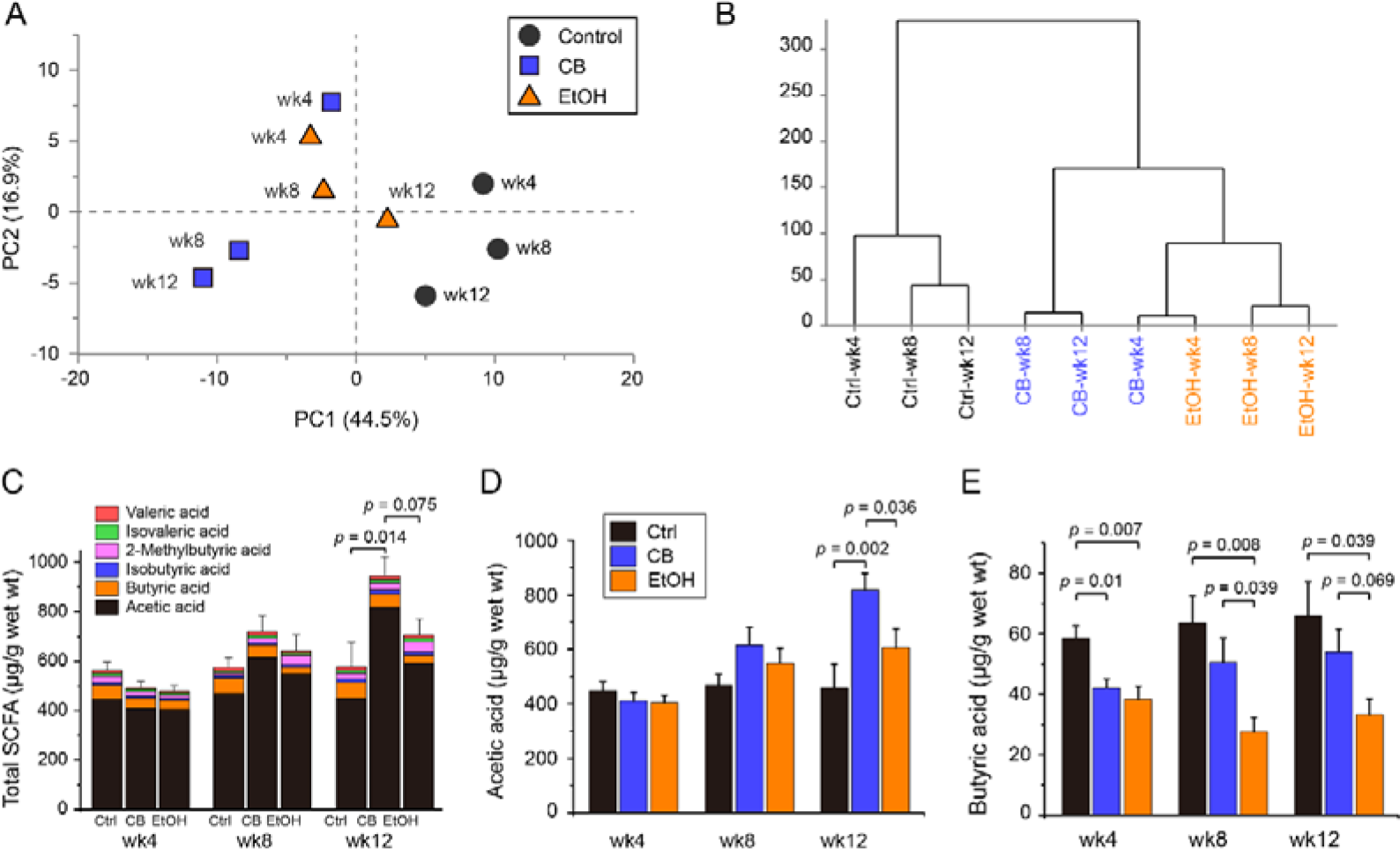
CB feeding and EtOH feeding caused difference in the caecal metabolome. (A) PCA scores plot showed the time-dependent “footprints” of the caecal contents metabolome. Each dot in the plots represents the mean value of the scores from the first and second principal components at a time point (n=8 mice/time point for the Ctrl group; n= 9-10 mice/time point for the experimental groups). (B) Clustering of gut metabolome based on mahalanobis distances calculated using ward linkage. (C) Total content of SCFAs in the colon (n=8 mice/time point for the Con group; n= 9-10 mice/time point for the experimental groups). Colon contents of (D) acetate and (E) butyrate. Bar plot is shown as mean ± SEM. Significance was evaluated using the two-tailed unpaired Student’s *t*-test.

### Correlations between key OTUs, differential metabolites and SCFAs

Understanding the interactions of microbiome and their metabolome holds promise for functional characterization of gut microbiota and for understanding human health and disease^30^. As such, we established the association between altered gut microbiota and their metabolism using correlation analyses and visualized interactions of microbiome and their metabolome by examining the caecal microbial composition and its metabolite profile (Fig. 6).

**Figure 6.**
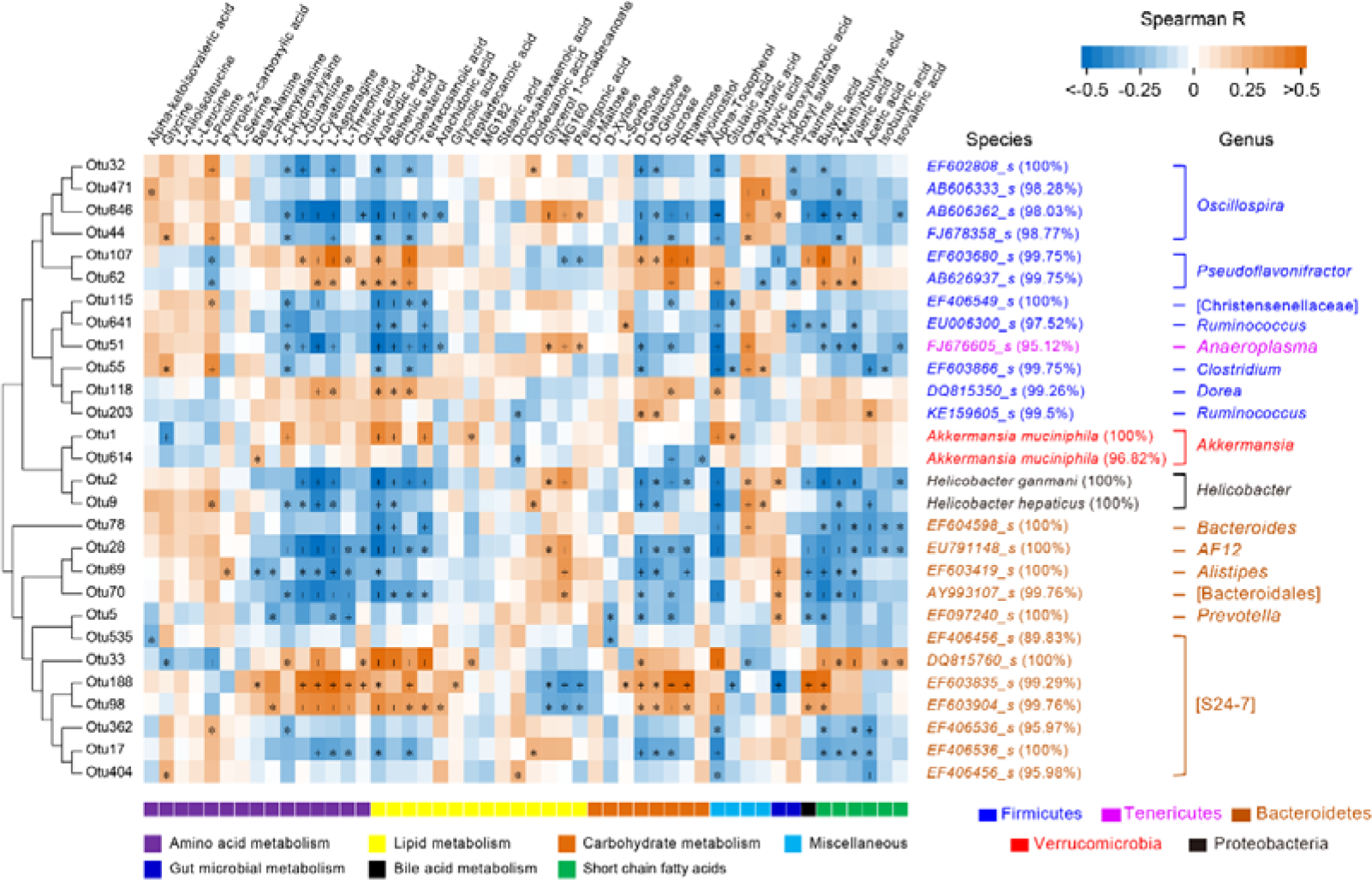
Heatmap shows the spearman correlations between key OTUs and differential metabolites and SCFAs. The color scale represents the spearman R value, with red and blue indicated positive and negative correlations, respectively. The metabolite types are denoted with the down color bar. Significance was evaluated using the two-tailed unpaired Student t-test. *p < 0.05; +p < 0.01.

At the OTU level, we observed a clustering of metabolites related to carbohydrate metabolism (rhamnose, galactose, glucose and sucrose), amino acid metabolism (L-cysteine, L-glutamine, quinic acid, 5-hydroxylysine) and lipid metabolism (arachidic acid, behenic acid, cholesterol and tetracosanoic acid) with a number of microbes, including a positive correlation between these metabolites and OTUs belonging to the genera *Pseudoflavonifractor, Dora, Ruminococcus, Akkermansia* and unclassified S24-7, and also a negative correlation was observed between these metabolites and OTUs belonging to the genera *Oscillospira, Anaeropiasma,* and *Helicobacter.* Interestingly, some metabolites related to lipid metabolism (dodecanoic acid, glycerol 1-octadecanoate and pelargonic acid) had inverse correlations with members of these genera indicating a diverse genetic potential of these OTUs. In addition, we also observed correlations between members of these genera with SCFAs, specifically a positive correlation with members of genera *Pseudoflavonifractor* and unclassified S24-7 and a negative correlation with the genera, *Oscillospira, Ruminococcus, Bacteroides, AF12, Alistipes, Prevotella* and unclassified S24-7.

### Low dosages of administered CB *vs.* EtOH were used to verify the hepatoprotective effect of compounds in CB

The above results suggested that the different interventions (CB *vs.* EtOH) led to differences in gut flora and their functions. These differences directly translated into differences in ALD with CB gavage resulting in lower levels of liver injury than gavage with EtOH. In order to verify whether a lower dosage of CB or EtOH affects liver following the same rules, we treated mice with a lower, 1.5-fold standard treatment incorporating either CB or EtOH (Fig. 7A). As expected, no obvious hepatic steatosis and lipid accumulation were observed in the CB-treated group during the entire experiment. However, light hepatic steatosis and lipid accumulation occurred in the EtOH-treated group at week 12 (Fig. 7B,C). Likewise, there were no significant differences in the level of plasma ALT and hepatic triglycerides between the CB-treated group and the control group. As for the EtOH-treated group, significantly increased levels of plasma ALT and hepatic triglycerides were observed from weeks 8 and 12 onward, respectively (Fig. 7D,E). These results further confirm our original hypothesis that chemical compounds in CB ameliorate ethanol-induced liver injury.

**Figure 7.**
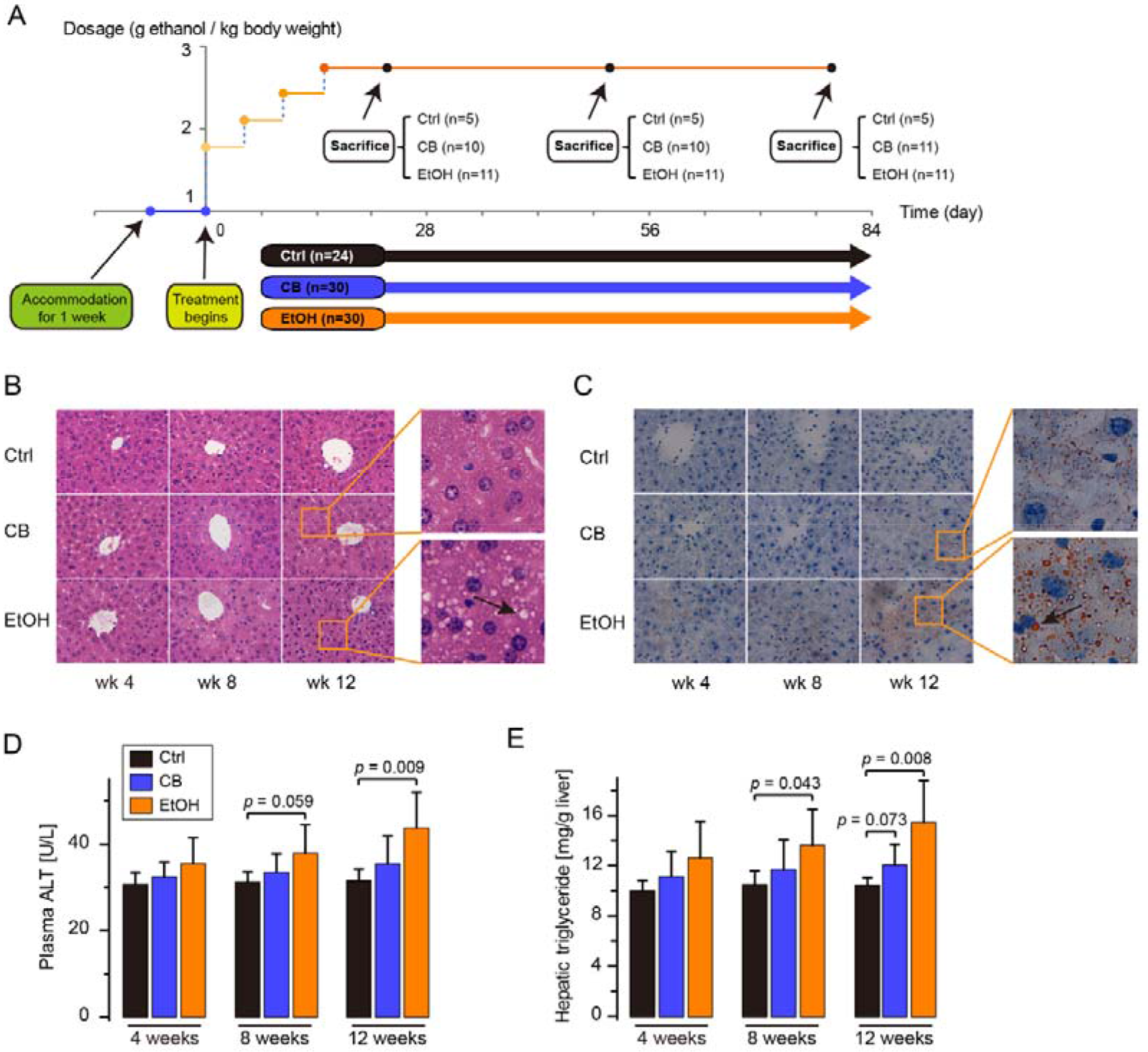
Effect of low dosage of Chinese Baijiu (CB) and ethanol (EtOH) intervention on liver. (A) Representative photomicrographs of H&E liver sections. (B) Representative photomicrographs of Oil Red O-stained liver sections. (C) Levels of plasma alanine aminotransferase (ALT). (D) Levels of hepatic triglyceride. Magnification, 400* (n=5 mice/time point for the Ctrl group; n=10-11 mice/time point for the experimental groups). Bar plot is shown as mean ± SEM. Significance was evaluated using the two-tailed unpaired Student *t*-test.

## Discussion

Baijiu is a type of traditional Chinese fermented alcoholic beverage, and also one of the earliest distilled liquors with written records that dated back to the Ming Dynasty (1368-1644 A.D.). Today, Baijiu is not only part of a unique cultural heritage, but has become a major consumer product with important economic values. Recent studies with the compositional analysis of CB determined that it contained abundant amounts of small molecule bioactive compounds^20, 31^. In this study, we hypothesized that these flavor compounds protect against ethanol-induced liver injury. To verify this hypothesis, we set out to compare the effect of CB and ethanol on liver through observation of how the gut microbiome and host metabolome were altered in response to ethanol and CB using a mouse model of chronic EtOH/CB gavage.

As expected, mice fed CB containing an equivalent amount of EtOH exhibited significantly lower levels of hepatic injury and steatosis than mice fed EtOH alone as indicated by biochemical and histological analyses. These results demonstrated that the compounds in CB partially offset the negative consequences of ethanol. Further microbiome analysis found that the abundance of caecal *Akkermansia muciniphila* was significantly higher in the CB group than that of EtOH group. Recent studies suggested that *Akkermansia* abundance was associated with the integrity of intestinal barrier function and that recovery of ethanol- or high-fat diet-induced *Akkermansia* depletion by oral administration of *Akkermansia* as a probiotic was effective in the restoration of intestinal barrier function and ameliorated both alcoholic and non-alcoholic liver disease^32, 33^. Dysfunction of the intestinal barrier is characterized by an increase in gut permeability which, in turn, leads to increased translocation of microbial products such as LPS, from the gut to the portal circulation resulting in elevated levels of circulating LPS^11, 34^ LPS can thus access the liver via the circulation and affect host metabolic physiology in critical ways. By activation of hepatic Kupffer cells and other recruited immune cells, LPS has been shown to induce the production of large amounts of pro-inflammatory cytokines resulting in liver inflammation and injury^34^. Our results indicated that CB treatment caused lower levels of liver injury. The flavor compounds contained in CB protected against ethanol-induced reduction of *Akkermansia*, therefore reducing gut permeability and translocation of pro-inflammatory LPS.

Two potential mechanisms reported for the observed higher abundance of *A. muciniphila* involved the antioxidant and antimicrobial effect of some compounds in CB. CB is abundant with many phenolic and heterocyclic compounds^20, 35^ which have been shown to possess remarkable antioxidant effects^36, 37^. *A. muciniphila* is an obligate anaerobe, lacking protection against free oxygen radicals^38^. Therefore, it is tempting to speculate that the oxygen radical scavenging capacity of these compounds can provide a survival advantage to *A. muciniphila.* Regarding the antimicrobial effect of these compounds, it has been suggested that phenolic compounds exert profound antimicrobial activity and are capable of stimulating the growth of *Akkermansia* in the transversal colon^39^. In addition, *Akkermansia* was found to be negatively associated with *Prevotella*^40^, whose relative abundance was significantly lower in the CB treatment group than in EtOH treatment group (Fig. S10). As such, the antimicrobial effect of CB could be associated with a reduction in the abundance of species capable of holding *Akkermansia* in check, thus favoring a rise in its proportion. Importantly, *Prevotella* has also been identified to be a LPS-producing genus with a pro-inflammatory function. In a mouse model of gut inflammation, animals colonized with *P. copri* had more severe disease than controls^41^. Whereas, *Akkermansia* was found to be a LPS-suppressing genus^42^. Significant lower relative abundance of the genera *Prevotella* and higher relative abundance of the genera *Akkermansia* in the CB treatment group could also explain why the liver and serum LPS level of the CB group were significantly lower than that seen in the EtOH group.

Another interesting finding from the current study was that there were significant differences observed in the caecal metabolome between CB-fed and EtOH-fed mice. These results indicated that ALD, like many other metabolic disorders such as nonalcoholic fatty liver disease^43^, type 2 diabetes^44^ and cardiovascular disease^45^, may be coordinately influenced by intestinal microbiome and metabolome.

Importantly, we observed a better metabolic activity of caecal microbiota of CB-fed mice that was manifested by elevated SCFA production, especially for acetate and butyrate. Similar to our findings, a previous study also observed that ethanol feeding increased the content of acetate in the gut lumen^13^. This can be explained by the enrichment of the functions related to ethanol metabolism of gut microbiota after ethanol feeding^46^. Acetic acid is the end product of ethanol metabolism. Higher concentrations of acetate in CB-fed mice suggested a higher detoxification for ethanol and/or acetaldehyde for the CB-fed mice^13^. As for butyrate, the other SCFA that was elevated in CB mice relative to EtOH mice, it plays multiple roles in intestinal and host health such as, enhancing the intestinal barrier function by activating AMP-activated protein kinase (AMPK) and regulating the assembly of tight junctions^47^. A recent study in experimental ALD found that butyrate supplementation protected against alcohol-mediated intestinal tight junction disruption and liver inflammation^48^. Butyrate is also an important energy source for intestinal epithelial cells and has been shown to be protective against colorectal cancer, partly by inhibiting histone deacetylases (HDACs)^49^ which are vital regulators of fundamental cellular events, such as cell cycle progression, differentiation, and tumorigenesis^50^. In addition to being an anti-tumor agent, butyrate is also a potent anti-inflammatory agent. By stimulating the development of regulatory T cells, butyrate can induce the production of the anti-inflammatory interleukin-10 (IL-10) which has the ability to prevent inflammatory reactions against pathogenic microbes^51, 52^ In fact, along with the cells that produce mucin (goblet cells) and those that maintain tight junctions (enterocytes), the gut immune cells are also considered part of the intestinal barrier^11^.

Another impressive finding concerning the metabolic differences between the two experimental groups CB and EtOH, was that CB-fed mice had higher levels of saturated LCFA (ie. dodecanoic acid, heptadecanoic acid, stearic acid, arachidic acid, behenic acid, tetracosanoic acid) than EtOH-fed mice (Fig. S4). Chronic ethanol administration has been reported to reduce the capacity of intestinal bacteria to synthesize saturated LCFA in mice and humans. Maintaining intestinal levels of saturated LCFA in mice improved eubiosis, stabilized the intestinal gut barrier, and reduced ethanol induced liver injury^24^. It restored eubiosis by preventing ethanol-induced overgrowth of intestinal bacterial and by increasing intestinal levels of probiotic *Lactobacillus*^24^, a bacteria with known beneficial effects in ALD, via multiple mechanisms, including maintaining gut barrier integrity^53, 54^ However, in our mice model, we did not find any OTU belonging to this genus in the caecum even in the control mice. This discrepancy might be due to several factors, such as different animal models of ALD (eg, *ad libitum vs.* intragastric ethanol administration), different durations of alcohol feeding, variability in the specific dietary components (eg, diets containing different types of fat and/or fermentable fibers), and also different sequencing methods (16S amplicon versus metagenome). Otherwise, it is possible that saturated LCFAs protect against ethanol-induced liver injury by other mechanisms than simply by increasing intestinal levels of probiotic *Lactobacillus.* Although the mechanism was not determined, we demonstrated that compounds in CB can prevent the loss of intestinal saturated LCFA and reduce ethanol-induced liver injury in mice. Our results support the concept that diets or agents designed to increase intestinal concentrations of saturated LCFA or their production by the intestinal microbiome might be developed to treat alcohol-induced liver disease^24^.

## Materials and methods

### Animal experiments

Specific Pathogen-Free (Spf) Wild-Type C57/B6 Mice (Male, Age 7-8 Weeks, Weight 20-23 G) Were Obtained From Shanghai Laboratory Animal Co, Ltd. (Slac, Shanghai, China). The Mice Were Housed In A Controlled Environment At 22 ± 2 °C With 40%-60% Humidity And A 12 H Light-Dark Cycle. Mice Had Access To Water And Fodder *Ad Libitum* And Their Body Weight And General Health Were Closely Monitored. After A One Week Acclimatization Period, Mice Were Randomly Divided Into Three Treatment Groups: The Control Group (Vehicle Only), Cb Group And Etoh Group. As It Was Not Practical To Enforce Each Mouse To Intake A Defined Dose Of Ethanol By *Ad Libitum*, All The Animals Were Given Etoh Or Cb Solutions By Oral Gavage. In Order To Be Consistent With The Experimental Groups, Normal Control Mice Received The Vehicle (Water). To Ensure The Consistent Gavage Volume, Cb And Etoh Were Diluted To The Same Etoh Concentration Before Use. The Initial Dosage Of Etoh (3.2 G/Kg Per Day) Was Increased In A Stepwise Manner 0.6 G/Kg Every 5 Days Until The 20Th Day (5.6 G/Kg Every) And Then Kept Constant To The End Of The Experiment. This Amount Of Etoh Was Equivalent To Approximately 3 Standard Drinks For A Human. The Confirmatory Animal Model Consisted Of Three Similar Groups: The Control Group, Cb Group And Etoh Group With A Lower Dosage Of The Treatments. The Initial Dosage Of Etoh (1.8 G/Kg Per Day) Was Increased In A Stepwise Manner 0.3 G/Kg Every 5 Days Until The 15Th Day (2.7 G/Kg Every) And Then Kept Constant Until The End Of The Experiment. The Amount Equivalent To About 1.5 Drinks In Humans. The Dosage Was Based On Standard Drink Amounts Set By Nih And Previous Work Published By Others^55^. Body Weight Gain And Food Intake Were Assessed Once A Week. Following 4, 8 Weeks Or 12 Weeks Of Cb Or Etoh Feeding, Mice Were Sacrificed By Cervical Dislocation After Fasting Overnight. Caecal Contents were excised and stored at -80°C immediately for further analysis. Blood was collected from the inferior vena cava, and citrated plasma was stored at -80°C for further analysis. Portions of liver and ileum tissues were fixed with 10% formalin for sectioning, while others were snap-frozen with liquid nitrogen and then stored at -80 °C. All mice were harvested between 08:00 and 11:00^56^ and received humane care in compliance with institutional guidelines.

### Biochemical analyses

Serum alanine aminotransferase (ALT) activity was colorimetrically measured using the Infinity ALT kit (Thermo Scientific). Hepatic triglyceride levels were measured using the Triglyceride Liquid Reagents Kit (Pointe Scientific, Canton, MI) according to the manufacturer’s instructions. Hepatic lipid peroxidation was quantified by measuring malondialdehyde using the thiobarbituric acid reactive substances (TBARS) assay (BioVision, Milpitas, CA, USA). Plasma LPS levels were measured by a commercial ELISA kit (CUSABIO, Wuhan, China).

### Staining Procedures

Formalin-fixed tissue samples were embedded in paraffin and stained with hematoxylin and eosin (H&E) to assess the histological features of steatosis and inflammation. For hepatic lipid accumulation analysis, frozen sections of fresh liver tissues embedded in OCT compound (Sakura) were cut and stained with Oil Red O. For immunofluorescence analysis of intestinal tight junction proteins, frozen intestinal sections were treated with anti-occludin and anti-claudin-2 antibodies (Invitrogen, Carlsbad, CA), followed by staining using a fluorescein isothiocyanate-conjugated secondary antibody (Servicebio, Wuhan, China). Laser-scanning confocal microscopy (Nikon D-Eclipse C1) was used for observation, and images were captured using a Nikon DS-U3 camera.

### Real-Time quantitative polymerase chain reaction

Total RNA from ileum tissue was isolated using TRIzol reagent (Invitrogen, Carlsbad, CA) according to the manufacturer’s instructions. The isolated RNA was then reverse transcribed with the TaqMan Reverse Transcription Reagents (Invitrogen) after assessment of RNA quantity. Semiquantitative analysis of relative gene expressions were performed on the Applied Biosystems 7500 Real Time PCR Systems (Applied Biosystems, Carlsbad, CA) using SYBR green PCR Master Mix (Qiagen). The sequences of the primers listed below 5’-CAACTGGTGGGCTACATCCTA-3’, Claudin-2 Forward 5’-CCCTTGGAAAAGCCAACCG-3’, Reverse 5’-TTGAAAGTCCACCTCCTTACAGA-3’, Occludin Forward 5 ’-CCGGATAAAAAGAGTACGCTGG-3 ‘), Reverse 5 ’-ACGGACC AGAGCGAAAGC AT-3 ‘, 18s rRNA Forward 5 ’-TGTCAATC CTGTCC GTGTCC-3 ‘, Reverse were designed from the Primer Bank (http://pga.mgh.harvard.edu/primerbank), and synthesized by Shanghai Sanggong Inc. (Shanghai, China). All primer pairs were validated by demonstrating high amplification efficiency, consistent single-peak melt curve, and the presence of single product of the expected amplicon size on agarose gels. The relative gene expression was normalized to 18s rRNA expression, and calculated by the 2^-ΔΔCT^ method^57^ setting the values of control group as one.

### Pyrosequencing of 16S rRNA gene V3-V4 region of gut microbiota

The metagenomic DNA of caecal samples used for pyrosequencing was extracted by a commercially available E.Z.N.A. Stool DNA Kit (Omega Bio-tek, Norcross, GA, U.S.) according to the manufacturer’s instructions. Hypervariable V3-V4 region of the prokaryotic 16S rRNA gene of each sample was amplified using a forward primer with a six-digit error-correcting barcode as described earlier^58^. After purification, the amplicons from each of the sample were equally combined to avoid or reduce PCR biases and were then subjected to a sequencing library preparation according to the manufacturer’s manual. DNA concentration and quality were determined by a NanoDrop 1000 spectrophotometer (NanoDrop Technologies, USA) and Qubit Fluorometer (Life Technologies, USA). Deep DNA pyrosequencing procedures were performed on a paired-end Illumina MiSeq PE300 (2 × 300 bp) platform at the Beijing Genomics Institute (BGI, Shenzhen, China) according to the manufacturer’s instructions.

### Metabolomic analysis of caecal contents by GC-TOFMS

The metabolites extraction and derivatization procedures for the caecal contents were described in previously published papers^59^ with minor modifications. One microliter of an aliquot of the derivatized solution was injected onto an Agilent 7890B gas chromatography coupled to a Pegasus HT time-of-flight mass spectrometer (Leco Corporation, St. Joseph, MI). Chemical compound separation was achieved on a DB-5MS capillary column (30 m x 250 μm I.D., 0.25-μm film thickness; (5%-phenyl)-methylpolysiloxane bonded and cross-linked; Agilent J&W Scientific, Folsom, CA). Helium was used as carrier gas at a constant flow rate of 1 mL/min. The temperature of injection, transfer interface, and ion source was set to 270, 260, and 200 °C, respectively. The GC temperature programming was set to 2 min of isothermal heating at 80 °C, followed by 10 °C/min oven temperature ramps to 180 °C, 6 °C/min to 230 °C, and 40 °C/min to 295 °C and a final 8 min maintenance at 295 °C. MS measurements were implemented with electron impact ionization (70 eV) in the full-scan mode (m/z 30-600), and the acquisition rate was 20 spectra/second in the TOFMS setting. All the samples were run in the order of “control-CB-EtOH-control”, alternately, to minimize systematic analytical deviations. A sample of mixed standards was used as quality control and was injected every 10 injections.

### Compound identification

The acquired data files from GC-TOFMS were processed by ChromaTOF software (v4.51.6.0, Leco, CA). After the pretreatment for baseline correction, denoising, smoothing, alignment, and deconvolution, raw data containing retention time, intensity, and the mass-to-charge ratio of each peak were obtained. Compounds were identified by comparing the mass fragments with NIST 05 standard mass spectral databases in NIST MS search 2.0 (NIST, Gaithersburg, MD) software with a similarity of >70% and finally verified by our in-house compound library (containing ~1000 mammalian metabolites). Both mass-spectrum and retention times were used to achieve precise compound annotations.

### Quantitative analysis of SCFAs

The extraction and derivatization procedures for SCFAs in caecal contents were described in published papers^60^. The GC-MS platform for quantitative analysis of SCFAs was the same as the one used for profiling caecal contents. The detection method had a minor modification. The temperature of injection, transfer interface, and ion source was set to 260, 290, and 230 °C, respectively. The GC temperature programming was set to 2 min of isothermal heating at 50 °C, followed by 10 °C/min oven temperature ramps to 70 °C, 3 °C/min to 85 °C, 5 °C/min to 110 °C, and 30 °C/min to 290 °C and a final 10 min maintenance at 295 °C. MS measurements were implemented with electron impact ionization (70 eV) in the full-scan mode (m/z 30-600) with an acquisition rate of 20 spectra/second in the TOFMS setting. All the samples were run in the order of “control-CB-EtOH-control”, alternately, to minimize systematic analytical deviations. A sample of mixed standards was used as quality control and was injected every 10 injections.

### Data analysis

For the microbiome data, quality control of the raw data was performed using BGI’s standard bioinformatics analysis pipeline. The acquired clean data with paired-end 16S rDNA reads were merged together using FLASH^61^. Operational Taxonomic Units (OTU) were clustered at 97% nucleotide similarity level using Uparse (version 7.0.1090)^62^ and aligned using RDP classifier (version 2.2) at 97% similarity level compared with the database of Greengene (version 201305)^63^. The alpha-diversity calculations including rarefactions and diversity indexes were performed using Mothur version 1.33.3^64^. Beta-diversity calculations of unweighted UniFrac principal coordinate analysis (PCoA) were performed using QIIME version 1.9.1^65^. Hierarchical cluster analysis (HCA) used for clustering of gut microbiota was calculated with multivariate analysis of variance (MANOVA). Redundancy analysis used for identifying Key OTUs responding to the CB and EtOH intervention was performed using CANOCO according to the manufacturer’s instructions^66^.

For the metabolome data, internal standards and any known artificial peaks, such as peaks caused by noise, column bleed, and the BSTFA derivatization procedure were removed from the data set. The resulting data were normalized to an internal standard prior to statistical analysis. The normalized data were mean centered and unit variance scaled during chemometric data analysis in the SIMCA-P+ software package (version 13.0, Umetrics, Umea, Sweden). Differential variables were selected with the criteria of variable importance in the projection (VIP > 1) in the PLS model and p < 0. 05 in a two-tailed unpaired *t*-test. The corresponding fold change shows how these selected differential metabolites varied between groups. To explore the functional correlation between the changes on metabolome perturbations and gut microbiome, Spearman’s correlation analyses were performed using SPSS v22 (IBM, USA).

### Data availability

The data that support the findings of this study are available from the authors on reasonable request. The mouse gut 16S rRNA gene sequencing data was deposited under NCBI BioProject PRJNA492383 and sequence reads are available at NCBI under BioSample IDs SAMN10101900.

## References

1. Organization WH, Unit WHOMoSA. Global status report on alcohol and health, 2014. World Health Organization (2014).

2. Forouzanfar MH, et al. Global, regional, and national comparative risk assessment of 79 behavioural, environmental and occupational, and metabolic risks or clusters of risks, 1990-2015: a systematic analysis for the Global Burden of Disease Study 2015. The Lancet 388, 1659–1724 (2016).

3. Rehm J, Samokhvalov AV, Shield KD. Global burden of alcoholic liver diseases. Journal of hepatology 59, 160–168 (2013).

4. Koppes L, Dekker J, Hendriks H, Bouter L, Heine R. Meta-analysis of the relationship between alcohol consumption and coronary heart disease and mortality in type 2 diabetic patients. Diabetologia 49, 648–652 (2006).

5. Koppes LL, Dekker JM, Hendriks HF, Bouter LM, Heine RJ. Moderate alcohol consumption lowers the risk of type 2 diabetes: a meta-analysis of prospective observational studies. Diabetes care 28, 719–725 (2005).

6. Mukamal KJ, et al. Moderate Alcohol Consumption and Chronic Disease: The Case for a Long-Term Trial. Alcoholism: Clinical and Experimental Research 40, 2283–2291 (2016).

7. Berg KM, et al. Association between alcohol consumption and both osteoporotic fracture and bone density. The American journal of medicine 121, 406–418 (2008).

8. Tilg H, Cani PD, Mayer EA. Gut microbiome and liver diseases. Gut 65, 2035–2044 (2016).

9. Schnabl B, Brenner DA. Interactions between the intestinal microbiome and liver diseases. Gastroenterology 146, 1513–1524 (2014).

10. Hartmann P, Seebauer CT, Schnabl B. Alcoholic liver disease: the gut microbiome and liver cross talk. Alcoholism: Clinical and Experimental Research 39, 763–775 (2015).

11. Szabo G. Gut-liver axis in alcoholic liver disease. Gastroenterology 148, 30–36 (2015).

12. Louvet A, Mathurin P. Alcoholic liver disease: mechanisms of injury and targeted treatment. Nature reviews Gastroenterology & hepatology, (2015).

13. Xie G, et al. Chronic ethanol consumption alters mammalian gastrointestinal content metabolites. Journal of proteome research 12, 3297–3306 (2013).

14. Nicholson JK, et al. Host-gut microbiota metabolic interactions. Science 336, 1262–1267 (2012).

15. Sharon G, Garg N, Debelius J, Knight R, Dorrestein Pieter C, Mazmanian Sarkis K. Specialized Metabolites from the Microbiome in Health and Disease. Cell Metabolism 20, 719–730 (2014).

16. Zheng X, et al. The footprints of gut microbial-mammalian co-metabolism. Journal of proteome research 10, 5512–5522 (2011).

17. Gu Y, et al. Analyses of gut microbiota and plasma bile acids enable stratification of patients for antidiabetic treatment. Nature Communications 8, 1785 (2017).

18. Kew W, Goodall I, Clarke D, Uhrin D. Chemical Diversity and Complexity of Scotch Whisky as Revealed by High-Resolution Mass Spectrometry. J Am Soc Mass Spectr 28, 200–213 (2017).

19. Schwarz M, Rodríguez M, Martínez C, Bosquet V, Guillén D, Barroso CG. Antioxidant activity of Brandy de Jerez and other aged distillates, and correlation with their polyphenolic content. Food Chemistry 116, 29–33 (2009).

20. Yao F, et al. Chemical analysis of the Chinese liquor Luzhoulaojiao by comprehensive two-dimensional gas chromatography/time-of-flight mass spectrometry. Sci Rep 5, 9553 (2015).

21. Wang L, et al. Intestinal REG3 Lectins Protect against Alcoholic Steatohepatitis by Reducing Mucosa-Associated Microbiota and Preventing Bacterial Translocation. Cell Host & Microbe 19, 227–239 (2016).

22. Ley RE, Turnbaugh PJ, Klein S, Gordon JI. Human gut microbes associated with obesity. Nature 444, 1022 (2006).

23. Yan AW, et al. Enteric dysbiosis associated with a mouse model of alcoholic liver disease. Hepatology 53, 96–105 (2011).

24. Chen P, et al. Supplementation of saturated long-chain fatty acids maintains intestinal eubiosis and reduces ethanol-induced liver injury in mice. Gastroenterology 148, 203–214. e216 (2015).

25. Grander C, et al. Recovery of ethanol-induced Akkermansia muciniphila depletion ameliorates alcoholic liver disease. Gut 67, 891–901 (2018).

26. Elamin EE, Masclee AA, Dekker J, Jonkers DM. Ethanol metabolism and its effects on the intestinal epithelial barrier. Nutr Rev 71, 483–499 (2013).

27. Muto S, et al. Claudin-2-deficient mice are defective in the leaky and cation-selective paracellular permeability properties of renal proximal tubules. Proc Natl Acad Sci U S A 107, 8011–8016 (2010).

28. Tripathi A, et al. The gut-liver axis and the intersection with the microbiome. Nature Reviews Gastroenterology & Hepatology, (2018).

29. Flint HJ, Duncan SH, Scott KP, Louis P. Links between diet, gut microbiota composition and gut metabolism. Proceedings of the Nutrition Society 74, 13–22 (2015).

30. Li M, et al. Symbiotic gut microbes modulate human metabolic phenotypes. Proceedings of the National Academy of Sciences 105, 2117–2122 (2008).

31. Yu T, et al. Overexpression of miR-429 impairs intestinal barrier function in diabetic mice by down-regulating occludin expression. Cell and Tissue Research 366, 341–352 (2016).

32. Grander C, et al. Recovery of ethanol-induced Akkermansia muciniphila depletion ameliorates alcoholic liver disease. Gut, (2017).

33. Plovier H, et al. A purified membrane protein from Akkermansia muciniphila or the pasteurized bacterium improves metabolism in obese and diabetic mice. Nat Med 23, 107–113 (2017).

34. Lucey MR, Mathurin P, Morgan TR. Alcoholic hepatitis. New England Journal of Medicine 360, 2758–2769 (2009).

35. Zhu S, et al. Characterization of flavor compounds in Chinese liquor Moutai by comprehensive two-dimensional gas chromatography/time-of-flight mass spectrometry. Anal Chim Acta 597, 340–348 (2007).

36. Yanagimoto K, Lee K-G, Ochi H, Shibamoto T. Anti oxidative activity of heterocyclic compounds found in coffee volatiles produced by Maillard reaction. J Agric Food Chem 50, 5480–5484 (2002).

37. Goldberg DM, Hoffman B, Yang J, Soleas GJ. Phenolic constituents, furans, and total antioxidant status of distilled spirits. J Agric Food Chem 47, 3978–3985 (1999).

38. Derrien M, Vaughan EE, Plugge CM, de Vos WM. Akkermansia muciniphila gen. nov., sp. nov., a human intestinal mucin-degrading bacterium. International journal of systematic and evolutionary microbiology 54, 1469–1476 (2004).

39. Kemperman RA, et al. Impact of polyphenols from black tea and red wine/grape juice on a gut model microbiome. Food Research International 53, 659–669 (2013).

40. Arumugam M, et al. Enterotypes of the human gut microbiome. nature 473, 174–180 (2011).

41. Scher JU, et al. Expansion of intestinal Prevotella copri correlates with enhanced susceptibility to arthritis. elife 2, e01202 (2013).

42. Jiang Z, Zhao M, Zhang H, Li Y, Liu M, Feng F. Antimicrobial Emulsifier-Glycerol Monolaurate Induces Metabolic Syndrome, Gut Microbiota Dysbiosis, and Systemic Low-Grade Inflammation in Low-Fat Diet Fed Mice. Mol Nutr Food Res 62, 1700547 (2018).

43. Dumas M-E, et al. Metabolic profiling reveals a contribution of gut microbiota to fatty liver phenotype in insulin-resistant mice. Proceedings of the national academy of sciences 103, 12511–12516 (2006).

44. Zhao L, et al. Gut bacteria selectively promoted by dietary fibers alleviate type 2 diabetes. Science 359, 1151–1156 (2018).

45. Wang Z, et al. Gut flora metabolism of phosphatidylcholine promotes cardiovascular disease. Nature 472, 57–63 (2011).

46. Dubinkina VB, et al. Links of gut microbiota composition with alcohol dependence syndrome and alcoholic liver disease. Microbiome 5, 141 (2017).

47. Peng L, Li Z-R, Green RS, Holzman IR, Lin J. Butyrate enhances the intestinal barrier by facilitating tight junction assembly via activation of AMP-activated protein kinase in Caco-2 cell monolayers. The Journal of nutrition 139, 1619–1625 (2009).

48. Cresci GA, Bush K, Nagy LE. Tributyrin Supplementation Protects Mice from Acute Ethanol-Induced Gut Injury. Alcoholism: Clinical and Experimental Research 38, 1489–1501 (2014).

49. Koh A, De Vadder F, Kovatcheva-Datchary P, Bäckhed F. From Dietary Fiber to Host Physiology: Short-Chain Fatty Acids as Key Bacterial Metabolites. Cell 165, 1332–1345 (2016).

50. Marks PA, Rifkind RA, Richon VM, Breslow R, Miller T, Kelly WK. Histone deacetylases and cancer: causes and therapies. Nature Reviews Cancer 1, 194 (2001).

51. Smith PM, et al. The microbial metabolites, short-chain fatty acids, regulate colonic Treg cell homeostasis. Science 341, 569–573 (2013).

52. Donaldson GP, Lee SM, Mazmanian SK. Gut biogeography of the bacterial microbiota. Nat Rev Micro 14, 20–32 (2016).

53. Wang Y, et al. Lactobacillus rhamnosus GG reduces hepatic TNFα production and inflammation in chronic alcohol-induced liver injury. The Journal of Nutritional Biochemistry 24, 1609–1615 (2013).

54. Wang Y, et al. Lactobacillus rhamnosus GG Treatment Potentiates Intestinal Hypoxia-Inducible Factor, Promotes Intestinal Integrity and Ameliorates Alcohol-Induced Liver Injury. The American Journal of Pathology 179, 2866–2875 (2011).

55. Wang D, et al. Green tea infusion protects against alcoholic liver injury by attenuating inflammation and regulating the PI3K/Akt/eNOS pathway in C57BL/6 mice. Food & Function 8, 3165–3177 (2017).

56. Zarrinpar A, Chaix A, Yooseph S, Panda S. Diet and Feeding Pattern Affect the Diurnal Dynamics of the Gut Microbiome. Cell Metabolism 20, 1006–1017 (2014).

57. Livak KJ, Schmittgen TD. Analysis of relative gene expression data using real-time quantitative PCR and the 2(T)(-Delta Delta C) method. Methods 25, 402–408 (2001).

58. Hamady M, Walker JJ, Harris JK, Gold NJ, Knight R. Error-correcting barcoded primers allow hundreds of samples to be pyrosequenced in multiplex. Nature methods 5, 235 (2008).

59. Gao X, Pujos-Guillot E, Sébédio J-L. Development of a quantitative metabolomic approach to study clinical human fecal water metabolome based on trimethylsilylation derivatization and GC/MS analysis. Analytical chemistry 82, 6447–6456 (2010).

60. Zheng X, et al. A targeted metabolomic protocol for short-chain fatty acids and branched-chain amino acids. Metabolomics: Official journal of the Metabolomic Society 9, 818–827 (2013).

61. Magoč T, Salzberg SL. FLASH: fast length adjustment of short reads to improve genome assemblies. Bioinformatics 27, 2957–2963 (2011).

62. Edgar RC. UPARSE: highly accurate OTU sequences from microbial amplicon reads. Nature methods 10, 996–998 (2013).

63. DeSantis TZ, et al. Greengenes, a chimera-checked 16S rRNA gene database and workbench compatible with ARB. Applied and environmental microbiology 72, 5069–5072 (2006).

64. Schloss PD, et al. Introducing mothur: open-source, platform-independent, community-supported software for describing and comparing microbial communities. Applied and environmental microbiology 75, 7537–7541 (2009).

65. Caporaso JG, Kuczynski J, Stombaugh J, Bittinger K, Bushman FD, Costello EK. QIIME allows analysis of high-throughput community sequencing data. Nat Methods 7, 335–336 (2010).

66. Ter Braak CJ, Smilauer P. CANOCO reference manual and CanoDraw for Windows user’s guide: software for canonical community ordination (version 4.5). Microcomputer Power: Ithaca, New York. (2002).

